# Transcriptional and neurochemical signatures of cerebral blood flow alterations in schizophrenia and the clinical high-risk state for psychosis

**DOI:** 10.1101/2024.03.13.583894

**Authors:** S.R. Knight, L. Abbasova, Y. Zeighami, J.Y. Hansen, D. Martins, F. Zelaya, O. Dipasquale, T. Liu, D. Shin, M.G. Bossong, M. Azis, M. Antoniades, O. Howes, I. Bonoldi, A. Egerton, P. Allen, O. O’Daly, P. McGuire, G. Modinos

**Author notes:** Shared first authorship (S.R. Knight, L. Abbasova).

## Abstract

The brain integrates multiple scales of description, from the level of cells and molecules to large-scale networks and behaviour, and understanding the relationships between these layers may be fundamental to advancing our understanding of how the brain works in health and disease. Recent neuroimaging research has shown that alterations in brain function that are associated with schizophrenia spectrum disorders (SSD) are already present in young adults at clinical high-risk for psychosis (CHR-P), yet the cellular and molecular determinants of these alterations are not well understood. Here, combining regional cerebral blood flow (rCBF) data with existing transcriptomic and neurotransmitter data, we show that cell-types involved in stress response and inflammation, as well as the dopamine, acetylcholine, GABAA and NMDA receptor systems, align as shared and distinct cellular and neurochemical signatures of rCBF phenotypes in people with SSD and those at CHR-P. Decoding the biological pathways involved in neuroimaging-based psychosis phenotypes may provide a basis for the development of novel interventions.

## INTRODUCTION

Schizophrenia spectrum disorders (SSD) are complex brain disorders with pharmacological treatments that are only partially effective, have many serious side-effects and are not preventative^1,2^. There is thus a major unmet need to develop new treatments for psychosis, and improving our understanding of its neurobiology is fundamental to achieving this goal.

Functional MRI (fMRI) is a widely available technique for mapping brain function safely and non-invasively. In recent years, MRI methods based on arterial spin labelling (ASL) perfusion contrast have been widely used to quantify regional cerebral blood flow (rCBF) in health and disease^3^. ASL provides a quantitative measurement of brain perfusion by magnetically labelling water as it enters the brain. This perfusion reflects an indirect measure of neural activity, as the metabolic demands of neural activity are served through the neurovascular coupling between neuronal activity and blood flow^4^. rCBF studies have demonstrated that subcortical hyperactivity and fronto-cortical hypoactivity are robust features of SSD^5^, proposed to underlie aberrant information processing and psychotic symptoms^6^. Subcortical rCBF hyperactivity is also present in people at clinical high-risk for psychosis (CHR-P)^5,7,8^. Moreover, hippocampal hyperactivity in CHR-P individuals is a predictor of poor functional outcome^9^, and has been found to normalise in those individuals who remit from the CHR-P state^10^, suggesting that these non-invasive measures may have potential for early detection and patient stratification. However, there is a lack of precise knowledge about how these measures are linked to dysfunction at cellular and molecular levels, which could be leveraged to guide new treatment strategies.

Recent open science initiatives now provide methods to integrate data across multiple scales, bridging macroscopic MRI-derived phenotypes with underlying microscopic cellular and molecular architecture. For instance, the Allen Human Brain Atlas (AHBA) is the most anatomically comprehensive gene expression dataset for the human brain, quantifying the expression of more than 20,000 genes by microarray^11^. *Imaging transcriptomics*^12^, which tests for covariance between the spatial expression pattern of each gene (AHBA) and the spatial topography of the neuroimaging pattern associated with the (clinical) phenotype, has been methodologically established^13–15^. This approach has led to a series of high-profile studies in psychosis^16–18^. For example, the map of structural MRI differences between people

with psychosis and healthy controls has been linked to the expression of genes that regulate synaptic signalling and nervous system development, including many genes implicated in the disorder by *post-mortem* studies^16^. Furthermore, new tools allow the integration of MRI phenotypes for a specific brain condition with existing maps of neurotransmitter binding and distribution (e.g. https://github.com/juryxy/JuSpace, https://github.com/netneurolab/neuromaps). Such multi-scale studies have demonstrated spatial covariance between drug-induced rCBF changes and *ex vivo* density of specific receptors^19^, and between antipsychotic-induced rCBF changes and D2/D3 receptor density derived from position emission tomography (PET)^20^. These *neuroreceptomic* analyses have also been applied to large-scale MRI datasets to uncover the neurochemical signatures of brain structural phenotypes^21–25^. For instance, in an analysis limited to the cortex, Hansen and colleagues examined the spatial correlation between 19 neurotransmitter systems and cortical thickness abnormalities in 13 different brain disorders, revealing that the serotonin transporter (5HTT) tracks the cortical thinning pattern of patients with SSD versus healthy controls^21^. How transcriptomic, cellular, and neurochemical attributes track rCBF alterations in SSD and CHR-P remains an open question.

Here, we bridged multiple brain spatial scales and data modalities to expose elements of microscopic architecture that may underlie rCBF abnormalities in individuals with SSD or at CHR-P. We integrated case-control rCBF maps of SSD and CHR-P with existing whole-brain microarray gene expression for 17,205 genes (AHBA) and whole-brain PET data from 19 receptors and transporters across nine neurotransmitter systems. We implemented an updated analysis pipeline using a functional brain atlas and incorporating subcortical brain regions to map the transcriptomic and neuroreceptor attributes of SSD and CHR-P-related rCBF patterns. We found that specific cell-type gene markers involved in stress response and inflammation track the patterns of rCBF alterations in SSD and CHR-P, as do the dopamine, GABA/glutamate, serotoninergic and cholinergic neurotransmitter systems. Collectively, these findings show how specific spatial patterns of gene expression and neurotransmitter systems overlap with rCBF alterations across the SSD spectrum, highlighting targets that may be responsive to intervention.

## RESULTS

### rCBF analysis

Individual rCBF data was acquired using ASL (see Supplement for MRI acquisition details and sample characteristics). rCBF data from 129 CHR-P individuals and 58 healthy controls (HC) were examined, comprising participants from two larger independent studies conducted at King’s College London, using the same ASL sequence^8,10^. rCBF data from 122 individuals with SSD and 116 HC were used from the multicentre CBFBIRN dataset, following standardised pre- processing analyses^26^. Unthresholded case-control t-stat maps (CHR-P vs HC; SSD vs HC) were derived from these data through independent samples t-tests in Statistical Parametric Mapping version 12 (SPM12; https://www.fil.ion.ucl.ac.uk/spm/). Of the 129 CHR-P individuals, 14 subsequently developed psychosis (CHR-T). For completeness, we further examined CHR-T vs CHR individuals who did not transition to psychosis (CHR-NT), as well as CHR-T vs HC and CHR-NT vs HC, reported in the Supplementary Materials. Finally, as an exploratory analysis, we also constructed case-control t-stat maps that included age, sex, and medication as covariates of no interest. Case-control rCBF t-stat maps were then parcellated into 100 cortical and 22 subcortical regions of interest (ROIs)^27,28^, with the mean value across each region z-scored to ease comparison across maps, resulting in a 122x1 vector of rCBF across the brain for each analysis (Fig. 1, Supplementary Fig. 1).

**Fig. 1.**
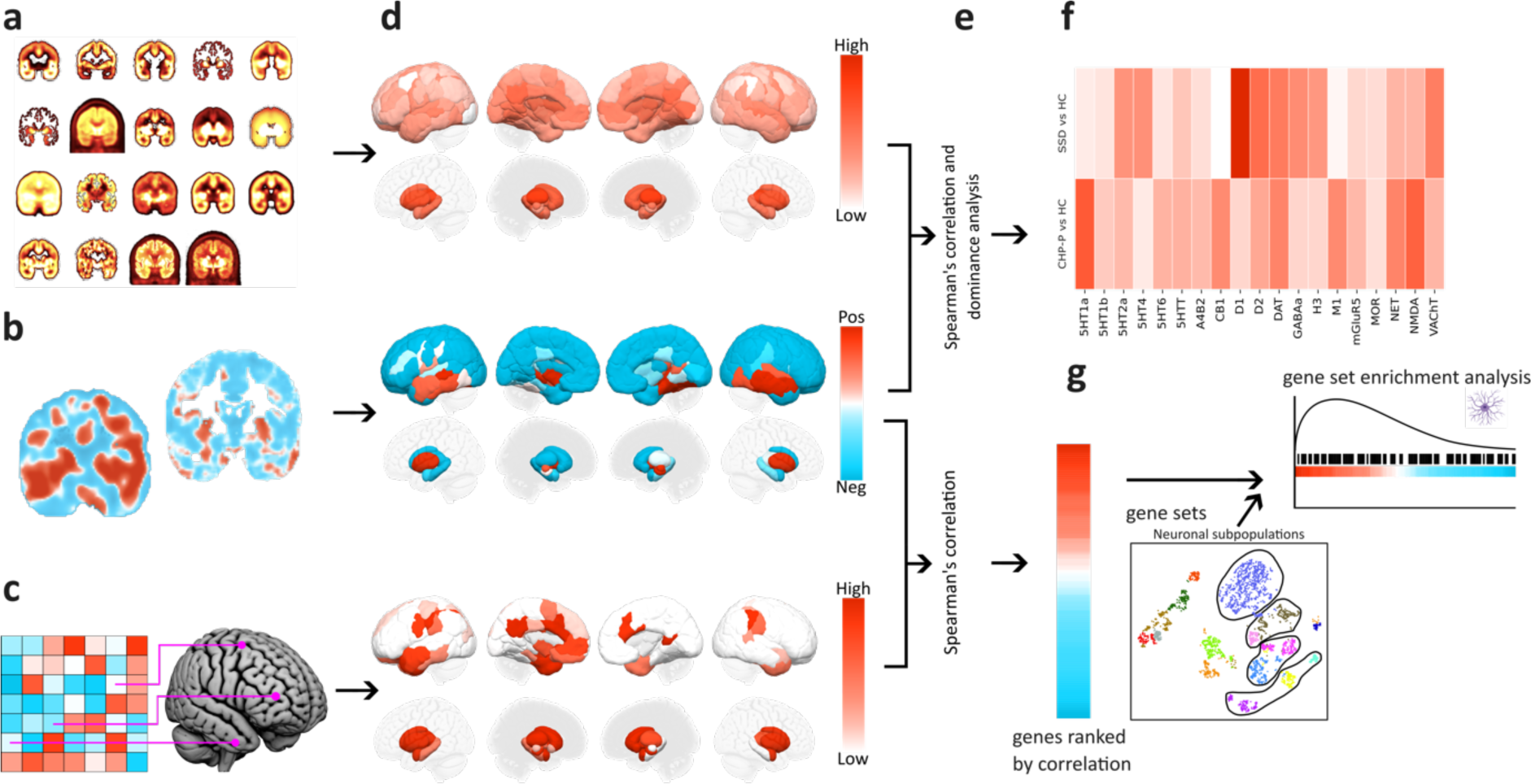
Analysis pipeline. **a,** 19 Group-level PET maps were obtained from http://github.com/netneurolab/hansen_receptors. **b,** Group-level case-control rCBF t-stat maps (SSD vs HC, CHR-P vs HC) were calculated in SPM12. **c,** Gene expression data was obtained from the AHBA. The AHBA tissue samples of donor brain images were registered to MNI space (ICBM152 nonlinear 2009c symmetric) using Advanced Normalisation Tools (https://zenodo.org/records/3677132). The gene expression data, comprising a total of 17,205 genes, were then pre-processed and log-normalised^29,30^. **d,** PET, transcriptomic and case-control rCBF maps were then parcellated into 122 regions of interest (ROI) using standardised brain atlases^27,28^, with the mean value extracted for each region. Three example distributions (NMDA, CHR-P vs HC, PTPRZ1) are displayed in the figure for illustrative purposes. The resulting averaged distributions within each of the 122 ROIs for each marker (PET map, case-control rCBF map and gene expression map) were z-scored and then subjected to **e,** Spearman’s correlation to rank all PET maps and genes according to their association with each rCBF marker (SSD vs HC, CHR vs HC, CHR-T vs CHR-NT), and **f,** dominance analysis to ascertain the unique contribution of each PET map to rCBF phenotypes (SSD vs HC, CHR-P vs HC, CHR-T vs CHR-NT). **g,** Finally, gene set enrichment analyses were conducted to ascertain specific cell-types involved, using marker genes obtained from a recently published single-cell transcriptomic study from the human brain^31^ and filtered by differential stability to retain genes with Pearson’s ρ > 0.5^29^. SSD, schizophrenia spectrum disorders. PET, positron emission tomography. CHR-P, clinical high-risk for psychosis. CHR-T, CHR-P individuals who subsequently transitioned to psychosis. CHR-NT, CHR-P individuals who had not transitioned at follow-up. HC, healthy control. MNI, Montreal Neurological Institute. rCBF, regional cerebral blood flow.

### Transcriptomic signatures of rCBF phenotypes

In both SSD and CHR-P, rCBF differences from HC significantly correlated with genes expressed in astrocytes, oligodendrocyte progenitor cells (OPCs), and vascular leptomeningeal cells (VLMCs) (Fig. 2b). CHR-P rCBF abnormalities were also associated with microglia and oligodendrocyte gene markers (Fig. 2b). In CHR-P, top correlating genes also fell into pathways involved in GPCR signalling and inflammation, as well as extracellular matrix (ECM) organisation. For SSD-associated rCBF differences, correlating genes were annotated to pathways involved in G protein-coupled receptor (GPCR) signalling, inflammatory responses, and metal ion binding (Fig. 2c). We performed the same analysis on the CHR-T rCBF phenotype but found no significant cell-type (Fig. 2b) or pathway (Fig. 2c) enrichments. This suggests that both similar and non-overlapping pathways are involved in SSD and CHR-P, but the CHR-P enrichments are not specific to those who subsequently transitioned to psychosis.

**Fig. 2.**
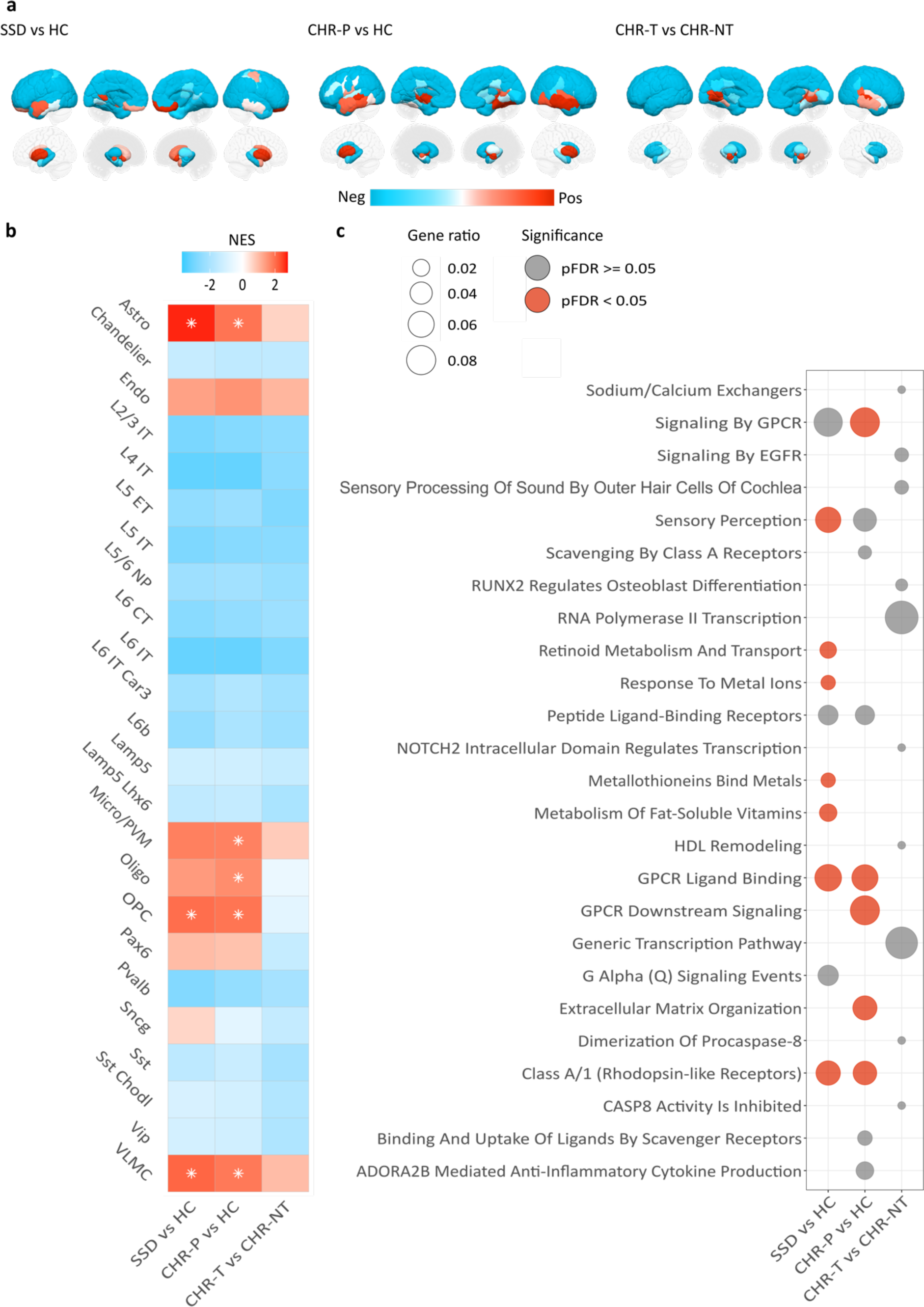
Transcriptomic contributions to SSD and CHR-P rCBF abnormalities. **a,** rCBF t-statistics parcellated by the 122-ROI Schaefer+Xiao atlas^27,28^, with the mean extracted for each region. **b,** Cell-type enrichments. Significant (PFDR < 0.05) enrichments are indicated with white asterisks and tiles are coloured by NES. **c,** Results of ORA analysis showing the union of the top 10 overrepresented Reactome pathway terms for each rCBF phenotype. Top pathways were selected by p-value. Significantly enriched (PFDR < 0.05) pathway terms are shown in red and point size represents the gene ratio, i.e. the number of genes falling into a particular pathway as a proportion of the total number of genes in the top 5% of positively correlated genes. NES, Net enrichment score. OPC, oligodendrocyte precursor cells. VLMC, vascular leptomeningeal cells.

Next, we sought to identify the cell-types expressing genes spatially correlated to the rCBF phenotypes using single-cell expression data and gene enrichment analyses (GSEA)^31^. Enrichment scores were adjusted for the number of genes for each given cell-type set to yield the normalised enrichment score (NES). We found that astrocyte, OPC, and VLMC gene modules correlated significantly (PFDR<0.0001) with both the SSD (NES of 2.66, 2.01, and 2.10, respectively) and CHR-P rCBF phenotypes (NES of 1.93, 1.93, and 1.87, respectively) (Fig. 2a). In addition, the CHR-P rCBF phenotype was significantly associated with oligodendrocytes (NES=1.62; PFDR<0.05) and microglia (NES=1.77; PFDR<0.0001). The CHR-T rCBF phenotype showed a similar correlation pattern with astrocytes, endothelial cells, microglia / PVM, and VLMCs, although these did not reach significance. These findings suggest that regions with greater expression of glial and vascular genes, which are presumably more abundant in the corresponding cell-types, show greater case-control differences in rCBF in the context of both SSD and CHR-P.

We next characterised whether the transcriptional signatures of the rCBF phenotypes of SSD and CHR-P capture clinically relevant information, such as sensitivity to a particular biological pathway, class of neurotransmitters, or specific signalling pathways. We performed overrepresentation analysis (ORA) on the top 5% of genes positively correlated with the SSD and CHR-P rCBF phenotypes to identify the enrichment of Reactome pathway terms^32,33^. The top SSD rCBF-linked genes showed significant enrichments for pathways relating to GPCR signalling in the contexts of both neurotransmission and immunity (involving genes such as *HTR1D, HTR2C, DRD3, CCL17, C3AR1, CXCL1, NTSR2*); sensory perception (including *TRIOBP, FSCN2, CDH23,* and multiple olfactory receptor genes); metal binding (multiple metallothioneins); and vitamin metabolism (including *SDC2, SDC4, LRP10, TTR, APOE, APOM, GPC5*) (Fig. 2b). The top CHR-P rCBF-linked genes were also enriched in genes involved in GPCR signalling, as well as ECM organisation (including *VCAM1, ICAM4, MMP28, COL1A2*, *SPARC*, *TIMP1, TIMP2*, *DCN*). Once again, the top CHR-T (vs CHR-P or HC) rCBF-linked genes were not significantly enriched in any pathway terms but still showed some association with genes involved in transcriptional regulation (predominantly comprising various transcription factors), regulation of caspase 8 (*CFLAR, RIPK1*), and to a lesser extent HDL remodelling (*LIPG*, *LCAT*) (Fig. 2b). The same analysis was repeated to test whether the top fraction of positively correlated genes were enriched in disease-related up- and down-regulated gene sets derived from a transcriptome-wide association study (TWAS) of schizophrenia, autism spectrum disorder, and bipolar disorder^34^. We found a significant (PFDR<0.0001) overrepresentation of genes upregulated in schizophrenia in SSD and CHR-P, but not CHR-T. We did not find enrichment for genes downregulated in schizophrenia.

### Neuroreceptor signatures of rCBF phenotypes

We next sought to examine whether specific receptor and transporter systems are linked to the patterns of rCBF abnormalities in our CHR-P and SSD samples. PET maps were parcellated into the 122 ROI atlas and the mean PET receptor/transporter density across each of these ROIs was then correlated with case-control rCBF t-stat maps through Spearman’s rank correlation. To assess significance, each correlation was compared against a null model using spatial-autocorrelation preserving surrogate maps (5,000 repetitions, pFDR<0.05)^35^. Negative correlations indicate areas where greatest case-control rCBF *hypofunction* is associated with higher receptor density, while positive correlations indicate areas of greater case-control rCBF *hyperfunction* and higher receptor density. The SSD-related rCBF phenotypes were negatively correlated with 5-HT2A and GABAA receptor densities (ρ=-.433, -.449, respectively, PFDR<0.05) (Fig 3a). Several other neurotransmitter systems also appeared positively correlated, such as D1, D2, DAT, and NMDA, but did not survive FDR correction. The CHR-P rCBF pattern of alteration was positively correlated with the distribution of D2 (ρ=.547, PFDR<0.05), DAT (ρ=.569, PFDR<0.05), NET (ρ=.319, PFDR<0.05) and NMDA receptors (ρ=.469, PFDR<0.05). This pattern was also positively correlated to the distribution of VAChT but did not survive FDR correction. Complementary analyses in the subset of CHR-P individuals who transitioned to psychosis vs those who did not revealed positive correlations between rCBF and DAT and NMDA receptor densities (ρ=.605, PFDR<0.05 and ρ=.526, PFDR<0.05 respectively). These findings support the notion that excitatory (NMDA) and inhibitory (GABAA) balance processes underlie brain functional abnormalities in SSD and CHR-P, as well as the involvement of the dopaminergic system in psychosis.

**Fig. 3.**
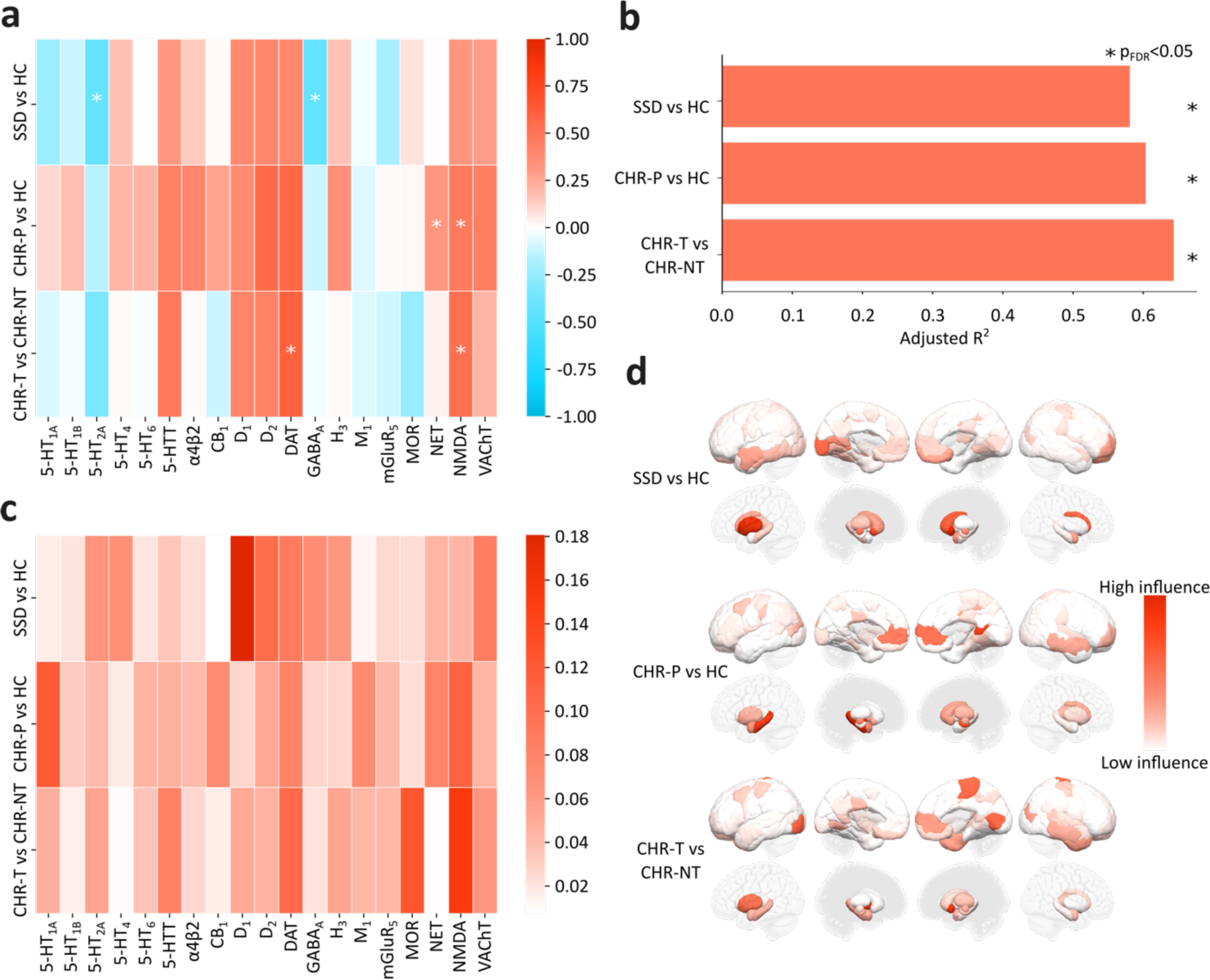
Neurochemical contributions to SSD and CHR-P rCBF abnormalities. **a,** Spearman’s correlations between parcellated rCBF and PET maps. Significant correlations, PFDR<0.05 marked *. **b,** Total dominance *R*^2^adj each case-control comparison**. c,** Dominance analysis of the unique contribution of receptors to case-control cerebral blood flow (rCBF) phenotypes. **d,** Cook’s distance estimates of most influential ROI in the receptor correlations. 5HT1A/1B/2A/4/6, serotonin 1A/1B/2A/4/6 receptor. 5HTT, serotonin transporter. α4β2, alpha-beta-4-nicotinic receptor. CB1, Cannabinoid receptor 1., D1/2, dopamine D1/2 receptor. DAT, dopamine transporter. GABAA, benzodiazepine binding site, γ-Aminobutyric acid A receptor. H3, histamine 3 receptor. M1, muscarinic receptor 1. mGluR5, Metabotropic glutamate receptor 5. MOR, μ-opioid receptor. NET, norepinephrine receptor. NMDA, *N*-methyl-D-aspartate receptor. VAChT, vesicular acetylcholine transporter.

To facilitate the interpretation of the above correlations given the high spatial collinearity between the different neurotransmitter systems (see^21^ for correlations between receptor maps), we conducted a multi-linear regression dominance analysis, where each predictor variable is iteratively entered into the model^36^. The resulting dominance of each predictor provides an intuitive grasp of the individual contribution of each receptor to the prediction of the rCBF differences map. The significance of the model fit was assessed against an FDR-corrected spatial autocorrelation-preserving null model^21,35,37^. Fig. 3c shows how the receptor maps covary with rCBF phenotypes. Receptor maps significantly predicted the rCBF phenotype of SSD (*R*^2^adj=.58, *P*FDR<.05) (Fig. 3d), with D1 (*R*^2^adj=.113[22%]), D2 (*R*^2^adj=.052[10%]), VAChT (*R*^2^adj=.049[10%]), and GABAA (*R*^2^adj=.042[8%]) as the most important predictors in the model. For individuals at CHR-P, receptor maps also predicted rCBF phenotype (*R*^2^adj=.6, *P*FDR<.05) (Fig. 3d), with 5HT1A (*R*^2^adj=.068[11%]), NMDA (*R*^2^adj=.062[10%]), DAT (*R*^2^adj=.048[8%]), NET (*R*^2^adj=.047[8%]), and M1 (*R*^2^adj=.047[8%]), and CB1 (*R*^2^adj=.046[8%]) as the most important predictors. When further dividing the CHR-P group based on subsequent psychosis transition (CHR-T vs CHR-NT), receptor maps significantly predicted the rCBF pattern (*R*^2^adj=.64, *P*FDR<.05). Unlike for CHR-P vs HC, MOR was the most important predictor of the CHR-T rCBF phenotype (*R*^2^adj=.083[15%]), followed by NMDA (*R*^2^adj=.081[14%]), DAT (*R*^2^adj=.05[9%]) and H3 (*R*^2^adj=.046[8%]). Taken together, these results demonstrate that overlapping and distinct molecular receptor distributions track regional alterations in brain function in SSD and CHR-P as measured non-invasively with ASL. More specifically, the dopaminergic system co-expresses with alterations in brain activity in both SSD and CHR-P, albeit to a stronger extent in SSD, and that this co-expression pattern may be preceded by involvement of the NMDA and other distinct neurotransmitter systems as suggested by both CHR-P and CHR-T.

### Sensitivity analyses

Repeating the above analyses including age, sex, and medication status as covariates of no interest rendered the transcriptomic relationships largely concordant with the original results (Supplementary Materials). This supports the specificity of associations between rCBF changes and astrocyte, microglia, oligodendrocyte, OPC and VLMC-associated gene markers across the extended psychosis spectrum, suggesting that these relationships were not simply driven by age, sex, or medication effects.

Neuroreceptor-rCBF correlations were relatively unchanged when including age and sex as covariates. However further inclusion of medication changed several correlational relationships (Supplementary Fig. 3). Briefly, the SSD-associated phenotype instead correlated with 5-HTT (ρ=.502, PFDR<0.05), D1 (ρ=.697, PFDR<0.05), DAT (ρ=.563, PFDR<0.05), and NMDA (ρ=.3996, PFDR<0.05). After including medication, only the CHR-P-associated phenotype’s correlation with NMDA survived FDR correction (ρ=.443, PFDR<0.05). The dominance analyses for SSD and CHR-P remained largely unchanged with the inclusion of age and sex as covariates, as well for the inclusion of medication for CHR-P. When medication was included as a covariate for the SSD-associated rCBF phenotype, the importance of D1 increased in the model. Together, these data suggest that the case-control rCBF by neuroreceptor correlations are in part related to medication effects, while the dominance analysis was robust to these effects.

To ensure parcellation method did not bias our results, we repeated the dominance analysis with the Desikan-Killiany atlas (82 ROIs). All models remained significant and the importance of individual receptors to the model was chiefly conserved between the two atlases (Supplementary Fig. 6).

Finally, we assessed where these rCBF-neuroreceptor associations were most strongly co-expressed in the brain. First, we assessed the influence of each ROI to our rCBF-neuroreceptor signatures through Cook’s distance analysis for each case-control comparison. SSD, CHR-P, and CHR-T rCBF-neuroreceptor correlations were most predominantly localised in subcortical regions, including the striatum, thalamus, and hippocampus (Fig. 3d, Supplementary Table 4). Supporting this subcortical predominance, when conducting the dominance analysis with only cortical regions (Schaefer 100 ROI cortical atlas), no models were significant (Supplementary Fig. 7). Together, these findings suggest that across the psychosis spectrum, rCBF-neuroreceptor associations are more closely linked in subcortical regions.

## DISCUSSION

In this study, multi-scale analyses integrating transcriptional, neurotransmitter and neuroimaging data reveal that rCBF alterations as measured noninvasively with ASL in CHR-P and SSD individuals are spatially coupled to distinct and overlapping patterns of whole-brain gene expression and neuroreceptor systems. More specifically, we found glial cell-type specific (i.e. astrocytes, oligodendrocytes, OPCs, and microglia) and VLMC markers to be most strongly correlated with case-control rCBF phenotypes across SSD and CHR-P, though not in CHR-T. Regarding neuroreceptor mapping, we found SSD, CHR-P, and CHR-T rCBF phenotypes to be related to the dopaminergic system. Moreover, GABAergic and NMDA receptors were differently linked to each disorder rCBF phenotype. Our findings also highlight further relatively less-established molecular determinants in SSD, CHR-P and CHR-T individuals. These findings provide new evidence for established theories that the cellular and molecular basis of rCBF phenotypes in psychosis relate to GABAergic/glutamatergic alterations^38–42^, abnormalities in dopaminergic signalling^9^, and white matter integrity^43^. With ASL being a technique known to be sensitive to drug-induced changes to specific neurotransmitter systems^19,20^, this technique may be particularly well suited to highlight potential biological targets for intervention.

Overall, our transcriptomic findings align with current hypotheses of psychosis pathophysiology^44–47^, and additionally highlight cell-types closely related to vasculature^48^. Consistent with the essential role of astrocytes in regulating rCBF and metabolic homeostasis^49^, these genes were found to be co-localised to more dysfunctional regions: those more likely to be metabolically active ‘hub regions’^50^. Astrocytes also play an important neurodevelopmental role in synapse formation and maintenance, hypothesised to underlie the pathophysiology of psychosis^51^ and other psychiatric disorders^52^. Interestingly, astrocytes are also important for the formation of perineuronal nets, as well as contributing to maintaining excitatory/inhibitory balance^53^, both proposed to be involved in the pathophysiology of psychosis^54^. Our correlations were robust for most sensitivity analyses, though did not survive FDR correction in our medication status sensitivity analysis in CHR-P or CHR-T.

Oligodendrocyte enrichment was also robustly positively correlated with the CHR-P phenotype, primarily in the subgroup that did not transition to psychosis (Supplementary Fig.2). Oligodendrocytes provide metabolic support for neurons, as well as myelinating synapses^55^. Multiple lines of evidence, including diffusion tensor imaging studies, have implicated white matter and myelination deficits across the psychosis spectrum^43,56,57^, and demyelination coincides with disease progression^58^. Moreover, histopathological and microscopy studies show oligodendroglia abnormalities reminiscent of apoptotic or necrotic cell death^59–61^. While the oligodendrocyte expression was not as robustly linked to the SSD phenotype as OPC expression, OPCs are tightly linked to oligodendrocyte differentiation^62^. Furthermore, we found evidence of microglia-specific gene markers correlating with rCBF phenotypes in CHR-P and SSD after including covariates, consistent with prior transcriptomic work implicating microglia in SSD^63^. Microglia play an essential role in mediating neuronal plasticity and synaptic pruning^64^, and microglial activation can be induced by psychological stress, brain injury or infection, so may represent a mechanism by which external stressors initiate or precipitate the development of SSD^65^. Microglial hyperactivation can have deleterious effects on other cell-types, including glial cells, thus potentially contributing to other cellular signatures of rCBF in SSD and CHR-P. Microglia also generate reactive oxygen species (ROS) in defence against deleterious factors, such as cellular debris and protein aggregates^66^. Oligodendrocytes and OPCs are especially sensitive to redox damage, due to glutathione depletion and high iron levels^67–69^. Pathological changes in oligodendrocytes and OPCs may in turn provoke microglial activation and astrocytic gliosis^70^, which can be attenuated through the administration of antipsychotics such as quetiapine and olanzapine^71,72^. Together, our findings fit with some previous imaging transcriptomic work co- localising gene expression with cortical thickness abnormalities in schizophrenia and other psychiatric disorders^17^, where positive correlations with both astrocyte- and microglia-specific genes were found.

We identified overrepresented pathways among genes correlating most strongly with rCBF phenotypes. In the SSD rCBF phenotype, top correlating genes were linked to pathways involved in GPCR signalling, inflammatory responses, retinoid metabolism, and metal ion binding. In the CHR-P phenotype, top correlating genes were associated with GPCR signalling and inflammation, as well as ECM organisation. We did not find significant pathway enrichments for the CHR-T rCBF phenotype. Previous work directly corroborating some of these genes and/or pathways in SSD has shown that, for example, levels of metallothionein MT-1 correlate negatively with psychotic symptom scores potentially serving as a potential biomarker for psychotic symptoms^73^. Retinoid dysregulation has also been implicated in schizophrenia^74,75^, and genes falling into enriched vitamin- / retinoid-linked metabolic pathways, such as retinoid transporters APOE and TTR, and ApoE receptor LRP10, are deregulated in schizophrenia^76–78^. Retinoic acid regulates the transcription of DRD2^79^ and other schizophrenia susceptibility genes, so a disruption in retinoic acid uptake and signalling may contribute to dopaminergic abnormalities observed in SSD. Many genes annotated to enriched pathways are also associated with cell-types and functions discussed above. For example, metallothioneins are enriched in astrocytes and are involved in binding and defence against oxidative damage^73^, and multiple genes involved in retinoid metabolism are more highly expressed in astrocytes, microglia and VLMCs. Our findings complement previous imaging transcriptomics work observing similar enrichment of genes upregulated in schizophrenia in regions with reduced morphometric similarity between cases and controls^16^. Overall, the correspondence between rCBF changes and spatial distributions of astrocytes, oligodendrocytes, OPCs, and microglia might arise due to both the increased vulnerability of oligodendrocytes and OPCs to neuroinflammation and oxidative stress, and/or their stimulation of immune responses due to intrinsic or extrinsic pathological abnormalities.

Regarding neuroreceptor mapping, we discovered prominent links between the distribution of specific neuroreceptors and SSD- and CHR-P-associated rCBF phenotypes. In SSD, dominance analysis showed that dopamine D1 receptor distribution was the strongest unique contributor to the rCBF phenotype, followed by D2 and DAT, consistent with the known role of the dopamine system in SSD pathophysiology^80^. The importance of these receptors and transporter systems in the dominance analysis remained after controlling for the effects of antipsychotic use. DAT was also an important predictor for the CHR-P and CHR-T rCBF phenotypes, even when including covariates. DAT is expressed in neurons that synthesise dopamine, and high DAT availability is considered a marker of excessive presynaptic dopamine activity^81^, which has been reported in CHR-P individuals by prior PET research^82,83^. Our results thus support a link between presynaptic dopaminergic abnormalities and case-control rCBF differences in CHR-P. In contrast to CHR-P vs HC, D1 was an important predictor of the CHR-T vs CHR-NT phenotype, as for SSD vs HC, which may point to a biomarker for transition to psychosis, although the number of participants in the CHR-T group was low (n=14).

Additionally, GABAA correlated with the SSD phenotype and was an important contributor in the dominance analysis. In turn, NMDA receptor distribution showed a significant positive correlation with the CHR-P vs HC rCBF phenotype, was an important predictor in the dominance analysis, survived all sensitivity analyses, and was also significant in the subset of CHR-P participants who subsequently transitioned to psychosis. Taken together, these results support the proposed role of glutamatergic (excitatory) and GABAergic (inhibitory) imbalance in SSD pathophysiology^84–87^. Interestingly, the observation of distinct (glutamatergic in CHR-P and GABAergic in SSD) and overlapping (dopaminergic) neuroreceptor contributions to rCBF phenotypes may be relevant to inform potential disease stage-specific targets.

Our analyses also revealed further, relatively novel, receptors/transporters linked to rCBF phenotypes. These included for example the cholinergic system in SSD: VAChT was an important predictor in our dominance model. Interestingly, acetylcholine is emerging as a promising new treatment target for both positive and negative symptoms in schizophrenia, as highlighted by a recent phase 3 trial indicating efficacy of the M1 and M4 agonist xanomeline-trospium^88^. M1 was among the most important predictors for CHR-P, but not SSD or CHR-T, phenotypes. We also found a negative association between 5-HT2A distribution and the SSD phenotype. *Post-mortem* prefrontal 5-HT2A reductions have been reported in SSD^89^, 5-HT2A agonists recapitulate some psychosis-relevant features^90^, and many atypical antipsychotics antagonise 5-HT2A^91^. Indeed, the correlations were no longer significant after including medication as a covariate. The complementary analysis of CHR-T vs CHR-NT individuals also revealed that the CHR-T phenotype strongly corresponded to MOR distribution. Interestingly, MOR has been linked to the first-episode psychosis multimodal neuroimaging phenotype in another study incorporating neurotransmitter systems, but not in chronic schizophrenia^92^. These data suggest MOR may be a novel target to redress rCBF dysfunction in prodromal psychosis. The additional associations observed with NET and 5- HT1A in the CHR-P group relative to the SSD findings were not unexpected, given the known heterogeneity in clinical outcomes in the CHR-P group^93^ and that these neurotransmitters have also been associated with depression, post-traumatic stress disorder, and generalised anxiety disorder^94–98^. Norepinephrine is well known to play a role in stress salience, as well as dopamine reuptake and excitatory/inhibitory signalling^99,100^. CB1 was also an important predictor of the CHR-P phenotype, which may be of relevance to cannabis use as a risk factor for psychosis and the antipsychotic potential of cannabinoids^101^. Finally, H3 was an important predictor for the CHR-T phenotype. H3 is known to have inhibitory effects on many neurotransmitter systems relevant to psychosis^102^ and H3’s therapeutic potential for treating cognitive dysfunction, including in psychosis, is currently being explored^103^. Altogether, our findings complement well-established and emerging theories of psychosis while highlighting further receptor systems that may be important for stratifying individuals at highest risk for psychosis and for follow-up investigation as potential therapeutic targets.

Overall, our neuroreceptomic findings are broadly consistent with prior research on receptor associations with SSD-related structural MRI phenotypes. Hansen et al.^21^ found that the strongest associations for the cortical thickness phenotype of SSD were with 5-HTT and D2 receptor distributions. While in our study both 5-HTT and D2 density correlated with SSD-associated rCBF, neither finding survived FDR correction. A possible explanation for this discrepancy might be that the cortical thickness phenotype of SSD reflects different processes, or that it reflects the consequences of long-term altered brain function on brain morphology. Indeed, rCBF alterations reflect current neuronal activity and have been proposed to precede and possibly drive subsequent structural abnormalities in SSD^38,104^. Furthermore, Hansen et al.’s analyses^21^ only included the 68 cortical regions in the Desikan– Killiany atlas used by the ENIGMA consortium^105^. For our rCBF analysis, we used a functional brain atlas comprising 100 cortical regions and 22 subcortical regions. Since subcortical abnormalities (including the striatum^106^ and hippocampus^42^) are among the most well-established findings in psychosis, our divergent findings may thus arise from including these SSD-relevant subcortical regions in our analyses. Cook’s distance analysis and the supplementary cortical-only dominance analysis confirmed that subcortical ROIs were among the most influential for rCBF-neuroreceptor co-expression. Future work characterising the cellular and molecular basis of functional and structural MRI phenotypes from the same individuals may expand our understanding of how multi-scale relationships vary or converge by imaging readout.

There are several limitations to note in our study. First, the benefit of using ASL-derived rCBF as a primary measure of brain function is that it is non-invasive, relatively inexpensive and, unlike BOLD fMRI, quantitative. However, its relatively low resolution and lower signal-to-noise ratio compared to structural MRI limits the granularity of the signal within ROIs. Whilst we demonstrate that the receptor analysis was consistent between different parcellation atlases, it nevertheless makes lower ROI count parcellation schemes more feasible. Second, not all SSD and CHR-P participants were medication naïve, and we were unable to control for treatment duration or medication dose (except for antipsychotics in SSD). Third, SSD and CHR-P data were from different datasets, acquired with different sequences and scanners, and initial CBF maps calculated with different methods, precluding a direct SSD versus CHR-P rCBF comparison. Fourth, both the transcriptomic (AHBA) and receptor atlases were constructed from independent groups of healthy volunteers of varying group sizes. Though not a limitation specific to our study, AHBA transcriptomic data were derived from six healthy post-mortem donors (of which five are male), and the cell-type markers used for GSEA were obtained from a single, separate study^31^. Though these datasets are cutting-edge, future work would benefit from more extensive datasets of gene expression. Regarding the receptor maps, the benefit of this technique is we can compare many PET maps, prohibitive in individual participants. However, as these maps were obtained from all over the world, in different participants and using different methodologies, we are limited to using normalised group-mean distribution rather than absolute values. Considering the hypothesis-generating nature of the present work, future research should follow up the observed correlations using direct experiments, for example experimental medicine studies involving pharmacological manipulation of 5-HT2A in SSD or MOR in CHR-P.

This work has significant scalability. Potential avenues for future work include a focus on other imaging modalities in the same people, such as diffusion-weighted tensor imaging, resting-state fMRI, or magnetoencephalography, which may provide a comprehensive account of the cellular and molecular underpinnings of different aspects of brain function as well as structural and functional connectivity. Ultimately, identifying the biological pathways underlying alterations that are observable by MRI, and determining the extent to which group phenotypes can be applied at the individual level, would allow us to understand the mechanisms that therapies can normalise in the specific people who show these MRI signatures, potentially providing a transformational change in mental health research.

In summary, by integrating available gene expression and neurotransmitter data with rCBF phenotypes in SSD and CHR-P, we provide novel evidence linking several specific cellular and molecular pathways to alterations in brain function in the psychosis spectrum. Our work provides a framework for a multi-scale understanding of the mechanisms underlying brain function and dysfunction in psychosis and provides new directions for unlocking the discovery of new pharmacological targets.

## METHODS

### Analysis of rCBF maps

Both SSD vs HC and CHR-P vs HC followed the same group analysis pipeline using the Statistical Parametric Mapping software (SPM12, [http://www.fil.ion.ucl.ac.uk/spm/software/spm12/]). Due to differences in scanning modality, pre-processing, and sample characteristics (see supplementary materials), each clinical group was compared to their own matched healthy control sample. Case-control t-stat maps of SSD vs HC and CHR vs HC were calculated using separate independent sample t-tests, including a grey matter mask (thresholded to contain voxels with a >20% probability of being grey matter) and covarying for global CBF. Additional maps were created for both SSD vs HC and CHR-P vs HC to include sex, age, and medication as covariates of no interest. To include medication as a covariate, 22 SSD individuals who did not have sufficient medication information were excluded. For SSD individuals, antipsychotic use was converted to chlorpromazine equivalent dose, and antidepressant use was categorised as binary values. For CHR individuals, antipsychotic, antidepressant, and anxiolytic medications were all categorised as binary values as dosages were not available for all participants. Information about anxiolytic medication use was not available in the SSD sample.

### Brain atlases

The case-control rCBF t-stat maps (unthresholded) and the PET receptor maps were parcellated into a combined atlas including 100 cortical^27^ and 22 subcortical^28^ ROIs. The Schaefer cortical atlas is constructed from a gradient-weighted Markov random field model, combining local and global similarity in resting-state fMRI gradients to parcellate the cortex. As the Schaefer atlas only covers cortical regions, this was combined with the 22 regions from the Xiao atlas, a high-resolution manual segmentation of subcortical regions, to create the combined Schaefer+Xiao atlas with 122 ROIs. The atlas was subsequently resampled to 2x2x2mm voxels using the nearest neighbour interpolation to match the resolution of the case-control rCBF maps.

### PET receptor maps

We used data and code made available at https://github.com/netneurolab/hansen_receptors for the PET receptor maps. These represent 19 different receptors and transporters, across 9 neurotransmitter systems^19,21,107–124^. As described by Hansen et al., all PET maps were acquired from healthy participants and each map was created as a group average across all participants within each study, resulting in a total of 1,235 participants (see Supplementary Table 2 or ref.^21^ for a complete list of tracers, ages, number of participants, and binding site). The receptor distributions of these maps have been previously validated through comparison with autoradiography data and alternative brain parcellations^21^. For our analyses relating such maps to an fMRI measure (rCBF), they were all parcellated into our 122 ROI atlas. The mean value across each region was z-scored to ease comparison across maps, resulting in a 122 x 19 matrix of relative receptor densities across the brain.

### Allen Human Brain Atlas (AHBA)

The AHBA dataset comprises microarray data of gene expression in post-mortem brain samples from six donors. See ref.^125^ and the Allen Human Brain Atlas website (https://human.brain-map.org/) for detailed information about donor demographics and the generation of this dataset. The AHBA tissue samples of donor brain images were registered to a MNI space template (ICBM152 nonlinear 2009c symmetric) using Advanced Normalisation Tools (ANTs, https://zenodo.org/records/3677132). The gene expression data, comprising a total of 17205 genes, were then pre-processed, log-normalised^29,30^. The results were then parcellated by our functional atlas, with the mean gene expression calculated across each of the 122 ROIs.

### Imaging transcriptomics analyses

rCBF t-stat maps and gene expression data, pre-processed as described above, were used as input to the imaging transcriptomics pipeline^126^ to calculate Spearman’s rank correlations between the mean expression of each gene and the rCBF case-control differences in each ROI. To determine the cell-types corresponding to any significantly correlated genes, gene set enrichment analysis (GSEA) was carried out using cell-specific gene markers derived from snRNA-seq of different regions of the human cerebral cortex. An unbiased, polygenic approach was adopted. Cell-type markers were obtained from snRNA-seq of different regions of the human cerebral cortex^31^. The resulting gene sets comprised markers of 16 transcriptionally defined cell subclasses, including eight interneuron subtypes (i.e. Pvalb, Sst, Sst Chodl, Lamp5, Lamp5 Lhx6, Sncg, Vip, Pax6), nine excitatory neuron subtypes (i.e., L2/3 IT, L4 IT, L5 IT, L6 IT, L6 IT Car3, L5 ET, L6 CP, L6b, L5/6 NP), and six non-neuronal subtypes (astrocytes, oligodendrocytes, OPCs, endothelial cells, VLMCs, and microglia/perivascular macrophages (PVMs); for more detail see Methods). Marker genes with low differential stability (Pearson < 0.5) were excluded to restrict analysis only to genes with reproducible patterns of differential expression^29^. Marker genes were filtered by log2 fold change to yield a maximum of 200 top genes per cell-type. As these markers reincluded recurring genes, corresponding to markers of the same cell-type across different cortical regions, duplicated genes were collapsed to produce unique gene sets for each cell-type. GSEA was carried out using the parcellated rCBF maps and spatial-autocorrelation preserving null maps to calculate FDR-corrected p-values for each cell-type enrichment score (see ‘Null models’). To identify pathway terms enriched among top deciles of positively correlated genes, ORA was performed using the fgsea package^32^ and Reactome 2022 pathway terms in Enrichr^33^, testing whether particular categories of genes are more likely to be enriched amongst the top rCBF- correlated gene set vs random background genes. We also performed this analysis using genes derived from a TWAS study of neuropsychiatric disorders^34^, separating genes significantly differentially expressed in schizophrenia (pFDR < 0.05) into up- (beta log2FC > 0) and downregulated (beta log2FC < 0) gene categories.

### Neuroreceptor mapping analyses

The unthresholded case-control rCBF t-stat maps and the 19 receptor map were parcellated by the Schaefer+Xiao 2x2x2mm atlas^27,28^ using the neuromaps toolbox in python^127^. Correlations between rCBF maps and receptor maps were assessed using Spearman’s correlations within neuromaps. To assess significance, 5000 nulls were generated for each comparison (see ‘Null Models’) and FDR-corrected for multiple comparisons, p<0.05. Cook’s distance analysis was conducted using statsmodels in python.

### Dominance analysis

Dominance analysis assesses the relative importance of a set of predictor variables in their contribution to the variance of a dependent variable^36^ (https://dominance-analysis.github.io/dominance-analysis/. This is achieved through iteratively fitting every combination of predictor variables into the same multiple linear regression model (2^p-1^, where p is the number of predictor variables). Predictor variables are compared in a pairwise fashion, in the context of models that contain a subset of all other predictors (2^p-2^), thus calculating the incremental R^2^ contribution of each predictor to the subset of all predictors. The total adjusted R^2^ of the model is the sum of these incremental contributions. Through a dominance analysis, the direct individual effect, importance to the total model, and importance relative to the other predictors can all be evaluated, making a dominance analysis intuitive and readily interpretable. Finally, each dominance analysis is normalised by the total fit of the model, for comparing between models. Our dominance analysis was conducted through code made available by Hansen et al.^21^ (https://github.com/netneurolab/hansen_receptors/), adapted to include our functional atlas of 122 ROIs including subcortical regions, and the ‘BrainSMASH’ generated null models (below).

### Null models

To account for spatial autocorrelation between the rCBF t-stat maps and PET and gene expression maps, both Spearman’s correlations and dominance analysis were tested against null models^37^. We implemented ‘BrainSMASH’ through the *netneurotools* package in Python (https://github.com/netneurolab/netneurotools). This method generates surrogate maps with autocorrelation that matches our input maps^35^, 5000 repetitions for each input map. To calculate a p-value, we counted the proportion of null models that performed better than the correlation/adjusted R^2^ of the alternative hypotheses.

### Data availability

The AHBA data are publicly available at https://human.brain-map.org/. Transcriptomic analysis was completed using code obtained from https://github.com/alegiac95/Imaging-transcriptomics. PET volumetric atlases and code were obtained and adapted from https://github.com/netneurolab/hansen_receptors and https://github.com/netneurolab/neuromaps. The case-control rCBF maps, other data, and code used in this paper will be made available upon publication.

### Author Contributions

SRK and GM conceived the study. SRK wrote the manuscript with LA, with valuable review and editing input from all authors. SRK and LA performed the formal analysis, with methodological contributions from SRK, YZ, JYH, and DM. SRK and LA interpreted the results. YZ, JYH, MB, AM, OH, IB, AG, AP, and PM provided data. GM provided supervision and secured funding. SRK was the project administrator.

