## Supplementary Materials for "Transcriptional and neurochemical signatures of cerebral blood flow alterations in schizophrenia and the clinical high-risk state for psychosis"

*Shared first authorship.

**Supplementary Methods**

**Individual rCBF maps and MRI acquisition**

*SSD vs HC*

The SSD sample comprised 122 individuals with a diagnosis of SSD and 116 HC as part of the CBFBIRN project.[1,2] Participants were recruited from seven study sites across the United States: Duke University, University of California, Irvine, University of California, San Francisco, University of Iowa, University of Minnesota, and the Mind Research Network. All participants provided written informed consent for the study. The protocol and consent forms were approved by the institutional review boards at each site. Imaging was performed on a 3T GE Excite System at Duke University and a Siemens Trio TIM at all other sites. ASL was performed using a standardised 2D single-shot FAIR protocol with pre-saturation pulses and QUIPSS II post-inversion saturation pulses. The ASL scan parameters were: TI1/TI2 = 600ms/1600ms, 10cm tag width, 1cm tag-slice gap, 220mm FOV, 24 slices (4mm thick, skip 1mm), TR 4 sec, 104 reps, spiral readout (TE = 3ms) for the GE system, partial Fourier EPI readout (TE = 12ms) for the Siemens system. Following the ASL scan, two 30-second scans were acquired with the ASL module turned off to obtain an estimate of the equilibrium magnetisation of cerebral spinal fluid and to correct for transmit/receive coil inhomogeneities. The raw CBF map was uploaded to the CBFBIRN pipeline[3] to generate individual CBF maps in physiological units of ml blood/100g tissue/min. As individual SSD and HC maps had already been processed by the CBFBIRN pipeline, we did not attempt to harmonise with CHR-P HC scans. An anatomical T1 scan was also acquired and registered to subjects' CBF space for segmentation of grey and white matter and for the generation of deformation fields to normalise the data into MNI152 for subsequent group analysis. Individual CBF maps were then pre-processed using the ASAP toolbox[4]. Individual CBF maps and structural T1 scans were reoriented into the same space, skull-stripped, and the skull-stripped T1 was then co-registered to the CBF map. Grey/white matter and CSF masks were created by segmenting the T1 image. The co-registered CBF map was then normalised to MNI space and smoothed with a 6mm kernel.

*CHR-P vs HC*

The CHR-P sample comprised 129 individuals meeting established CHR-P criteria (Yung et al. 2005), including 14 individuals who transitioned to psychosis during follow-up, and 58 HC as part of two previous studies conducted at King's College London[5,6]. Both datasets were acquired using the same General Electric Signa HDX 3.0T scanner at the Centre for

Neuroimaging Sciences, the same ASL sequence, and participants were assessed using the same clinical methods. Full recruitment details and MRI acquisition methods are described in detail in the original publications[5,6]. Ethical approval for both studies was obtained from the National Health Service UK Research Ethics Committee, and all participants provided written informed consent to participate in the study. Individual CBF maps were pre-processed using the same pipeline (ASAP toolbox) as the SSD and HC scans described above.

*Sensitivity analysis*

To ensure our results were not due to atlas choice, we repeated the analysis with the 83 ROI Desikan-Killiany atlas[7], resampled to ASL space using FSL. We subsequently removed the brainstem ROI, resulting in an 82 ROI atlas with 68 cortical and 14 subcortical ROIs. To confirm the importance of subcortical regions in our analysis, we repeated the analysis with only the 100 cortical ROIs of the Schaefer atlas[8], resampled to ASL space.

**Supplementary Table 1. Sample characteristics.** HC, healthy control group. CHR-P, Clinical high-risk for psychosis, CHR-T, clinical high risk for psychosis who subsequently transitioned to psychosis. CAARMS, Comprehensive Assessment of At-Risk Mental States. PANSS, Positive and Negative Syndrome Scale.

| | | CBFBIRN | | NEUTOP/PROD | | |
|---|---|---|---|---|---|---|
| | | HC (n = 116) | SSD (n = 122) | HC (n= 58) | CHR-P (n= 129) | CHR-T (n =14) |
| **Age (mean, S.D.)** | | 36.86 (11.12) | 39.51 (11.55) | 24.77 (4.3) | 22.66 (4.09) | 22.25 (4.21) |
| **Sex (*n*, M, F)** | | 83, 34 | 92, 30 | 32, 26 | 73, 56 | 8, 6 |
| **Race (%)** | **White** | 80% | 64% | 64% | 50% | 57% |
| | **Black** | 7% | 16% | 2% | 19% | 29% |
| | **Asian** | 9% | 14% | 3% | <1% | 7% |
| | **Other/unspecified** | 4% | 6% | 31% | 30% | 7% |
| **Years in education (mean, S.D.)** | | 15.4 (1.77) | 13.4 (1.9) | 14.11(2.51) | 13.6 (2.54) | 14.55(2.76) |
| **Handedness (% Right)** | | 96% | 88% | 86% | 81% | 100% |
| **CAARMS** | **Pos** | - | - | - | 8.28 | 9.64 |
| | **Neg** | - | - | - | 5.56 | 14 |
| **PANSS** | **Pos** | - | 12.13 | - | 13.88 | 15.77 |
| | **Neg** | - | 12.6 | - | 12.88 | 14.08 |
| **Antipsychotic use (*n*, chlorpromazine equivalent dose)** | | - | 101 (386.25mg) | - | 17 | 1 |
| **Antidepressant use (*n*)** | | - | 40 | - | 35 | 2 |
| **Antianxiety use (*n*)** | | - | - | - | 1 | 1 |
| **Cannabis use, lifetime (%)** | | 90 (78%) | 96 (79%) | 27 (47%) | 66 (51%) | 5 (36%) |
| **Global rCBF (ml/100g/min, mean, S.D)** | | 42.04(15.6) | 42.77 (11.31) | 43.86(9.27) | 44.39(9.59) | 42.67 (10.65) |

### Supplementary analysis of CHR-P phenotypes

For completeness we repeated our analyses with the subgroups of CHR-P participants that subsequently transitioned to psychosis (CHR-T) to HC participants, as well as CHR-P who did not transition (CHR-NT) with HC.

*Transcriptomic signatures*

In CHR-T vs HC, replicating our results for CHR-T vs CHR-NT, we did not find any significant correlating genes or pathway enrichments (Supplementary Fig 2., $P_{FDR}$>0.05). Comparing CHR-NT vs HC, rCBF differences correlated with genes expressed in astrocytes, oligodendrocytes, oligodendrocyte progenitor cells (OPCs), Pax6, and vascular leptomeningeal cells (VLMCs)

(Supplementary Fig. 2). Correlating genes were annotated to pathways involved in G protein-coupled receptor (GPCR) signalling.

*Neuroreceptor signatures*

The correlations found in CHR-T vs CHR-NT were even stronger when comparing CHR-T vs HC ($\rho$=.685, P<0.05 and $\rho$=.589, $P_{FDR}$<0.05 respectively), while interestingly no correlations reached significance for CHR-NT vs HC (Supplementary Fig. 3). The dominance analysis of CHR-T vs HC was significant and resembled the analysis of CHR-T vs CHR-NT ($R^2_{adj}$=.68, $P_{FDR}$<.05), with NMDA ($R^2_{adj}$=.11[16%]), DAT ($R^2_{adj}$=.08[11%]), and MOR ($R^2_{adj}$=.06[9%]) as the most important predictors. The dominance analysis of CHR-NT vs HC was also significant ($R^2_{adj}$=.49, $P_{FDR}$<.05) and resembled analysis of the CHR-P vs HC phenotype, but with 5-HT1a ($R^2_{adj}$=.1[20%]), NET ($R^2_{adj}$=.05[10%]), CB1 ($R^2_{adj}$=.05[10%]), and 5-HT6 ($R^2_{adj}$=.04[9%]) being more important predictors than a de-emphasised NMDA ($R^2_{adj}$=.04[7%]) (Supplementary Fig. 5).

**Sensitivity analysis of covariates**

**Transcriptomic signatures sensitivity analysis**

Controlling for effects of age + sex produced an additional enrichment for endothelial cells in CHR-P (NES=1.58; $P_{FDR}$<0.05) (Supplementary Fig. 2a). Regressing out effects of age + sex + medication resulted in the loss of astrocyte enrichments but retention of endothelial enrichments. Oligodendrocyte, OPC, microglia, and VLMC enrichments also remained significant when covariates were included. The SSD rCBF phenotypes gained significant correlation with oligodendrocytes (NES=1.37, $P_{FDR}$<0.05) and microglia (NES=1.70, $P_{FDR}$<0.0001) upon regression of age and sex, but the association with oligodendrocytes and VLMCs lost significance when controlling for medication status (Supplementary Fig. 2a). Nevertheless, enrichment for OPCs remained significant across the tested conditions. The patterns of enrichment broadly held after controlling for covariates. However, no significantly overrepresented pathway terms were detected in CHR-P when controlling for age + sex, and in CHR-T (Supplementary Fig 2b). The sensitivity analysis supports the specificity of the association between rCBF changes and OPC-associated gene markers across the extended psychosis spectrum. It suggests that this relationship is not simply driven by age, sex, or medication effects.

**Neuroreceptor signatures sensitivity analysis**

The SSD rCBF-related correlations with 5-$HT_{2A}$ and $GABA_A$ receptor densities remained significant when including sex and age as covariates (all $p_{FDR}$<0.05, Supplementary Fig. 3). However, further adding antidepressant and antipsychotic medication as covariates in the model rendered the 5-$HT_{2A}$ and $GABA_A$ correlations non-significant. Instead, we found correlations with 5-HTT (ρ=.502, $P_{FDR}$<0.05), $D_1$ (ρ=.697, $P_{FDR}$<0.05), DAT (ρ=.563, $P_{FDR}$<0.05), and NMDA (ρ=.3996, $P_{FDR}$<0.05). The CHR-P rCBF-related correlations with the distribution of $D_2$, DAT, NET and NMDA receptors remained significant after including age and sex as covariates (all $P_{FDR}$<0.05, Suppl. Fig. 3), and a new positive correlation was found with VAChT (ρ=.495, $P_{FDR}$<0.05). However, after additionally including antidepressant, antipsychotic, and anxiolytic medication (one participant) as covariates, only the association with NMDA survived FDR correction (ρ=.443, $P_{FDR}$<0.03). Finally, we tested the potential effects of confounders on the dominance analyses. The pattern involving important contributions of $D_1$, $D_2$, and VAChT receptor densities to the prediction of the SSD-related rCBF phenotype remained unchanged after including age and sex as covariates ($R^2_{adj}$=.58, $P_{FDR}$<.05) (Supplementary Fig. 5). Additional inclusion of medication as a covariate further improved the model ($R^2_{adj}$=.63, $P_{FDR}$<. 05) and increased the dominance of $D_1$ (from $R^2_{adj}$=.1[18%] to $R^2_{adj}$=.16[25%]). In terms of the CHR-P related rCBF phenotype, the model implicating DAT, $M_1$, 5-$HT_{1A}$, $CB_1$, NET, and NMDA receptors remained significant after controlling for age and sex, and additionally medication, with similar unique contribution of receptors ($R^2_{adj}$=.63, $P_{FDR}$<0.05 and $R^2_{adj}$=.55, $P_{FDR}$<0.05 respectively) (Supplementary Fig. 5).

**Supplementary Table 2. Receptors and transporters in neuroreceptor analysis.** BPND, non-displaceable binding potential; VT, tracer distribution volume; Bmax, density (pmol ml−1) converted from binding potential (5-HT) or distributional volume (GABA) using autoradiography-derived densities; SUVR, standard uptake value ratio. Values in parentheses indicates number of females. * indicates transporter. See Hansen et al.[9] for additional details.

| Receptor/transporter | Neurotransmitter | Tracer | Measure | *n* | Age (mean, S.D.) | Reference |
|---|---|---|---|---|---|---|
| 5-HT1A | Serotonin | [11C]WAY-100635 | $BP_{ND}$ | 35 (17) | 26.3 ± 5.2 | Savli et al.[10] |
| 5-HT1B | Serotonin | [11C]P943 | $BP_{ND}$ | 23 (8) | 28.7 ± 7.0 | Savli et al.[10] |
| 5-HT2A | Serotonin | [11C]Cimbi-36 | $B_{max}$ | 29 (14) | 22.6 ± 2.7 | Beliveau et al.[11] |
| 5-HT4 | Serotonin | [11C]SB207145 | $B_{max}$ | 59 (18) | 25.9 ± 5.3 | Beliveau et al.[11] |

| | | | | | | |
|---|---|---|---|---|---|---|
| 5-HT6 | Serotonin | $[^{11}C]$GSK215083 | $BP_{ND}$ | 30 (0) | 36.6 ± 9.0 | Radhakrishnan et al.[12] |
| 5-HTT* | Serotonin | $[^{11}C]$DASB | $B_{max}$ | 100 (71) | 25.1 ± 5.8 | Beliveau et al.[11] |
| α4β2 | Acetylcholine | $[^{18}F]$Flubatine | $V_T$ | 30 (10) | 33.5 ± 10.7 | Hillmer et al.[13] |
| CB1 | Cannabinoid | $[^{11}C]$OMAR | $V_T$ | 77 (28) | 30.0 ± 8.9 | Normandin et al.[14] |
| D1 | Dopamine | $[^{11}C]$SCH23390 | $BP_{ND}$ | 13 (7) | 33 ± 13 | Kaller et al.[15] |
| D2 | Dopamine | $[^{11}C]$FLB-457 | $BP_{ND}$ | 55 (29) | 32.5 ± 9.7 | Sandiego et al.[16] |
| DAT* | Dopamine | $[^{123}I]$-FP-CIT | SUVR | 174 (65) | 61 ± 11 | Dukart et al.[17] |
| GABA | GABA | $[^{11}C]$Flumazenil | $B_{max}$ | 16 (9) | 26.6 ± 8 | Nørgaard et al.[18] |
| H3 | Histamine | $[^{11}C]$GSK189254 | $V_T$ | 8 (1) | 31.7 ± 9.0 | Gallezot et al.[19] |
| M1 | Acetylcholine | $[^{11}C]$LSN3172176 | $BP_{ND}$ | 24 (11) | 40.5 ± 11.7 | Naganawa et al.[20] |
| mGluR5 | Glutamate | $[^{11}C]$ABP688 | $BP_{ND}$ | 73 (48) | 19.9 ± 3.04 | Smart et al.[21] |
| MOR | Opioid | $[^{11}C]$Carfentanil | $BP_{ND}$ | 204 (72) | 32.3 ± 10.8 | Kantonen et al.[22] |
| NET | Acetylcholine | $[^{11}C]$MRB | $BP_{ND}$ | 77 (27) | 33.4 ± 9.2 | Ding et al.[23] |
| NMDA | Glutamate | $[^{18}F]$GE-179 | $V_T$ | 29 (8) | 40.9 ± 12.7 | Galovic et al.[24] |
| VAChT* | Acetylcholine | $[^{18}F]$FEOBV | SUVR | 18 (13) | 66.8 ± 6.8 | Aghourian et al.[25] |

### Case-control rCBF t-stat maps

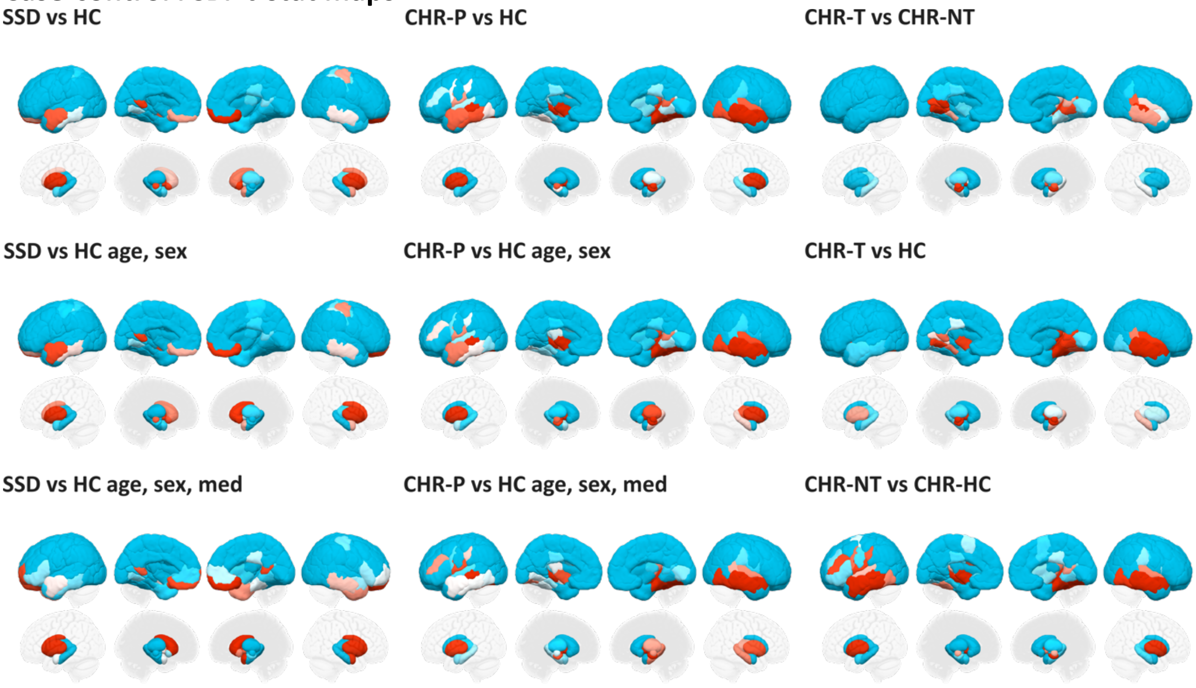

**Supplementary Fig. 1.** rCBF case-control t-stat maps (unthresholded)

### Transcriptomic analysis

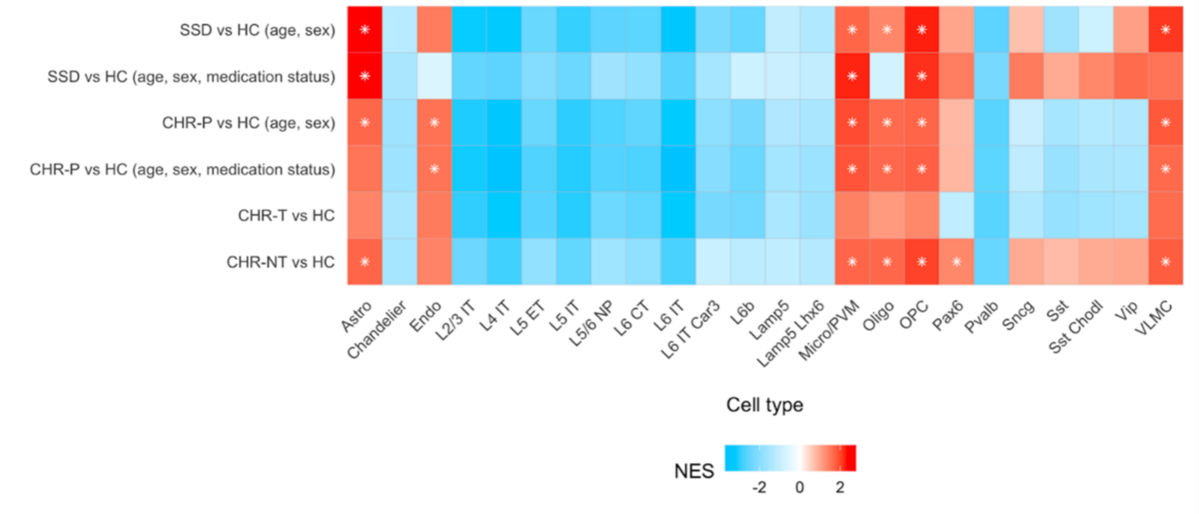

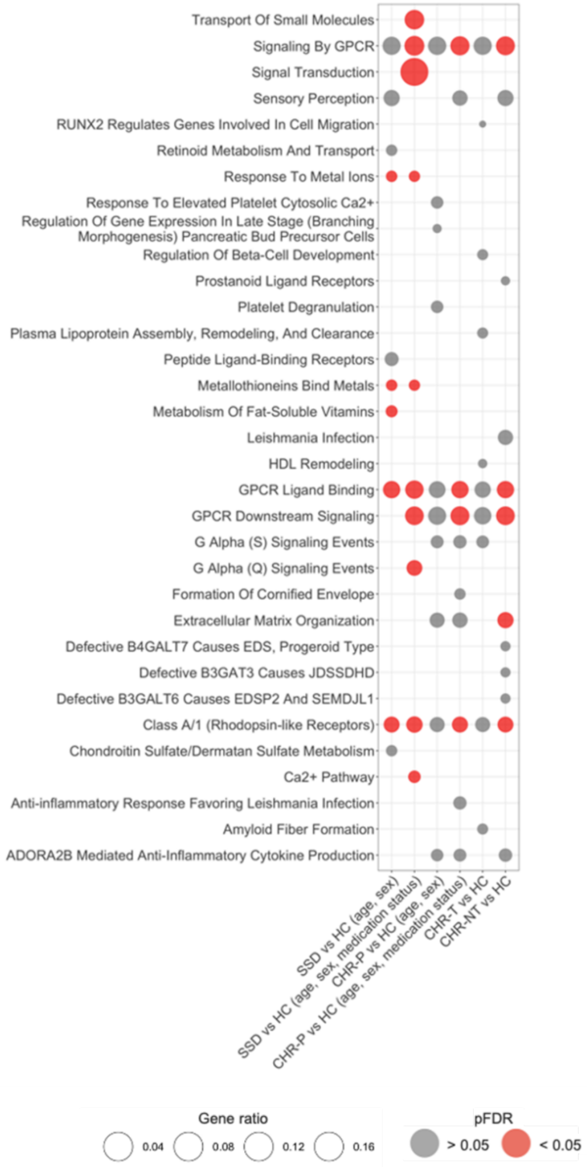

**Supplementary Fig. 2. Results of imaging transcriptomics analysis.** SSD vs HC and CHR-P vs HC sensitivity analysis controlling for covariates (age, sex, medication status). Also showing results for CHR-T vs HC and CHR-NT vs HC. (2a) Cell-type enrichments. Significant (pFDR < 0.05) enrichments are indicated with white asterisks and tiles are coloured by NES. (2b) Results of ORA analysis showing the union of the top 10 overrepresented Reactome pathway terms for each rCBF phenotype. Top pathways were selected by p-value. Significantly enriched ($p_{FDR}$<0.05) pathway terms are shown in red and point size represents the gene ratio.

### Neuroreceptomic analysis

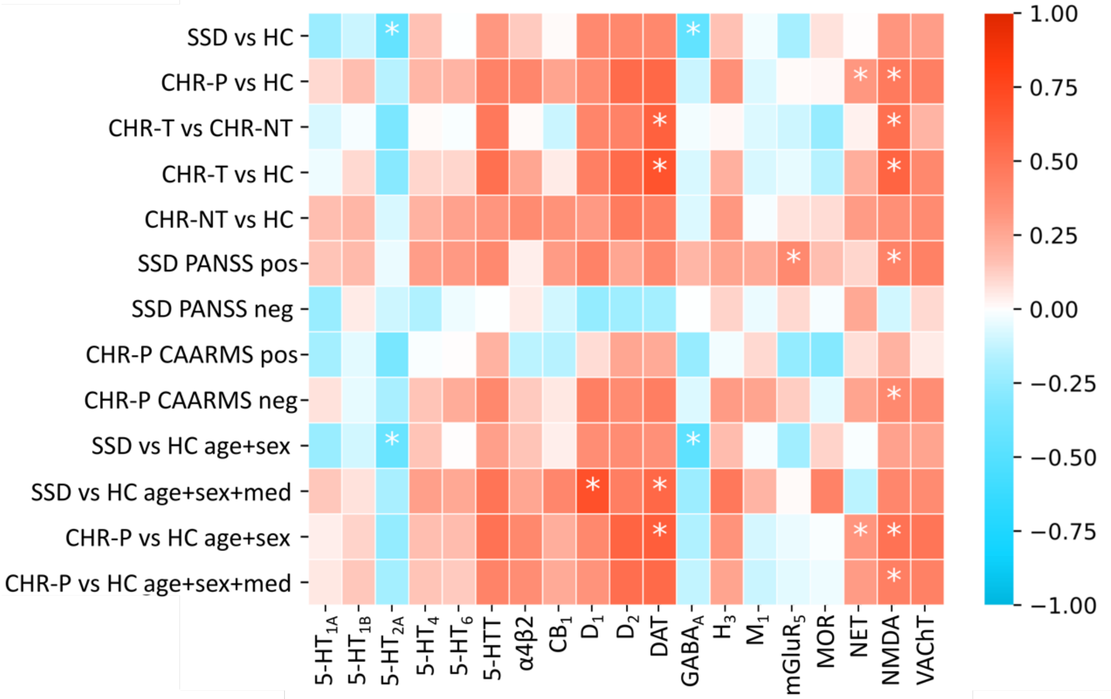

**Supplementary Fig. 3.** Spearman's Correlations between parcellated rCBF and PET maps. Significant correlations, $P_{FDR}<0.05$ marked *. 5HT1a/1b/2a/4/6, serotonin a/1b/2a/4/6 receptor. 5HTT, serotonin transporter. A4B2, alpha-beta-4-nicotinic receptor. CB1, Cannabinoid receptor 1. D1/2, dopamine D1/2 receptor. DAT, dopamine transporter. GABAa, benzodiazepine binding site, γ-Aminobutyric acid A receptor. H3, histamine 3 receptor. M1, muscarinic receptor 1. mGluR5, Metabotropic glutamate receptor 5. MOR, µ-opioid receptor. NET, norepinephrine receptor. NMDA, *N*-methyl-D-aspartate receptor. VAChT, vesicular acetylcholine transporter.

**Supplementary Table 3. Spearman's correlations**

| | 5HT1a | 5HT1b | 5HT2a | 5HT4 | 5HT6 | 5HTT | A4B2 | CB1 | D1 | D2 | DAT | GABAa | H3 | M1 | mGluR5 | MU | NET | NMDA | VAChT |
|---|---|---|---|---|---|---|---|---|---|---|---|---|---|---|---|---|---|---|---|
| **SSD vs HC** | -0.234 | -0.117 | -0.433 | 0.157 | -0.005 | 0.317 | 0.138 | 0.014 | 0.39 | 0.397 | 0.397 | -0.449* | 0.162 | -0.029 | -0.196 | 0.076 | 0.003 | 0.323 | 0.295 |
| **CHR-P vs HC** | 0.094 | 0.171 | -0.152 | 0.198 | 0.206 | 0.426 | 0.396 | 0.272 | 0.378 | 0.547 | 0.569 | -0.113 | 0.345 | -0.078 | 0.009 | 0.017 | 0.319* | 0.469* | 0.442 |
| **CHR-T vs CHR-NT** | -0.081 | -0.02 | -0.332 | 0.014 | -0.013 | 0.484 | 0.013 | -0.114 | 0.411 | 0.424 | 0.605* | -0.029 | 0.016 | -0.074 | -0.108 | -0.25 | 0.034 | 0.526* | 0.21 |

*Cook's distance*

**Supplementary Table 4.** Cook's distance analysis displaying 20 most influential ROIs (of 122) for each case-control comparison, ordered by descending influence.

| SSD vs HC | | CHR-P vs HC | | CHR-T vs CHR-NT | |
|---|---|---|---|---|---|
| Right nucleus accumbens | 0.2304 | Left subthalamic nucleus | 0.1640 | Left globus pallidus externa | 0.4264 |
| Left putamen | 0.1402 | Left hippocampus | 0.0930 | Left subthalamic nucleus | 0.0967 |
| Left nucleus accumbens | 0.0690 | 7Networks_RH_Default_PCC_1 | 0.0769 | Left globus pallidus interna | 0.0951 |
| Right caudate | 0.0616 | Right substantia nigra | 0.0707 | Right nucleus accumbens | 0.0871 |
| 7Networks_LH_Vis_5 | 0.0572 | 7Networks_RH_Default_PFCm_1 | 0.0525 | 7Networks_RH_Vis_5 | 0.0585 |
| Left globus pallidus externa | 0.0522 | Right subthalamic nucleus | 0.0478 | 7Networks_LH_Vis_4 | 0.0585 |
| Left red nucleus | 0.0512 | 7Networks_LH_Default_PFC_3 | 0.0476 | 7Networks_RH_SomMot_8 | 0.0565 |
| Right red nucleus | 0.0390 | Left red nucleus | 0.0405 | Left putamen | 0.0562 |
| 7Networks_LH_Vis_6 | 0.0379 | Left amygdala | 0.0374 | 7Networks_RH_Default_PFCm_1 | 0.0318 |
| Right globus pallidus externa | 0.0336 | Left globus pallidus externa | 0.0290 | 7Networks_RH_Limbic_TempPole_1 | 0.0292 |
| Left caudate | 0.0295 | Right nucleus accumbens | 0.0276 | 7Networks_RH_Vis_7 | 0.0286 |
| Right amygdala | 0.0292 | Right caudate | 0.0270 | Left hippocampus | 0.0249 |
| 7Networks_LH_SalVentAttn_Med_2 | 0.0286 | Left globus pallidus interna | 0.0267 | Left substantia nigra | 0.0242 |
| 7Networks_RH_Cont_PFCl_1 | 0.0285 | 7Networks_RH_SalVentAttn_Med_1 | 0.0267 | Right red nucleus | 0.0239 |
| 7Networks_LH_Default_Temp_1 | 0.0278 | Right red nucleus | 0.0255 | 7Networks_RH_Default_Temp_1 | 0.0236 |
| 7Networks_RH_Limbic_OFC_1 | 0.0276 | 7Networks_RH_Default_Temp_2 | 0.0253 | 7Networks_LH_Vis_3 | 0.0224 |
| 7Networks_LH_Default_PCC_1 | 0.0263 | 7Networks_RH_Default_Temp_1 | 0.0247 | Left amygdala | 0.0221 |
| 7Networks_LH_Limbic_TempPole_2 | 0.0195 | Left putamen | 0.0246 | Right subthalamic nucleus | 0.0211 |
| 7Networks_LH_Vis_3 | 0.0182 | Right thalamus | 0.0219 | 7Networks_LH_Cont_Cing_1 | 0.0192 |

*Dominance analysis*

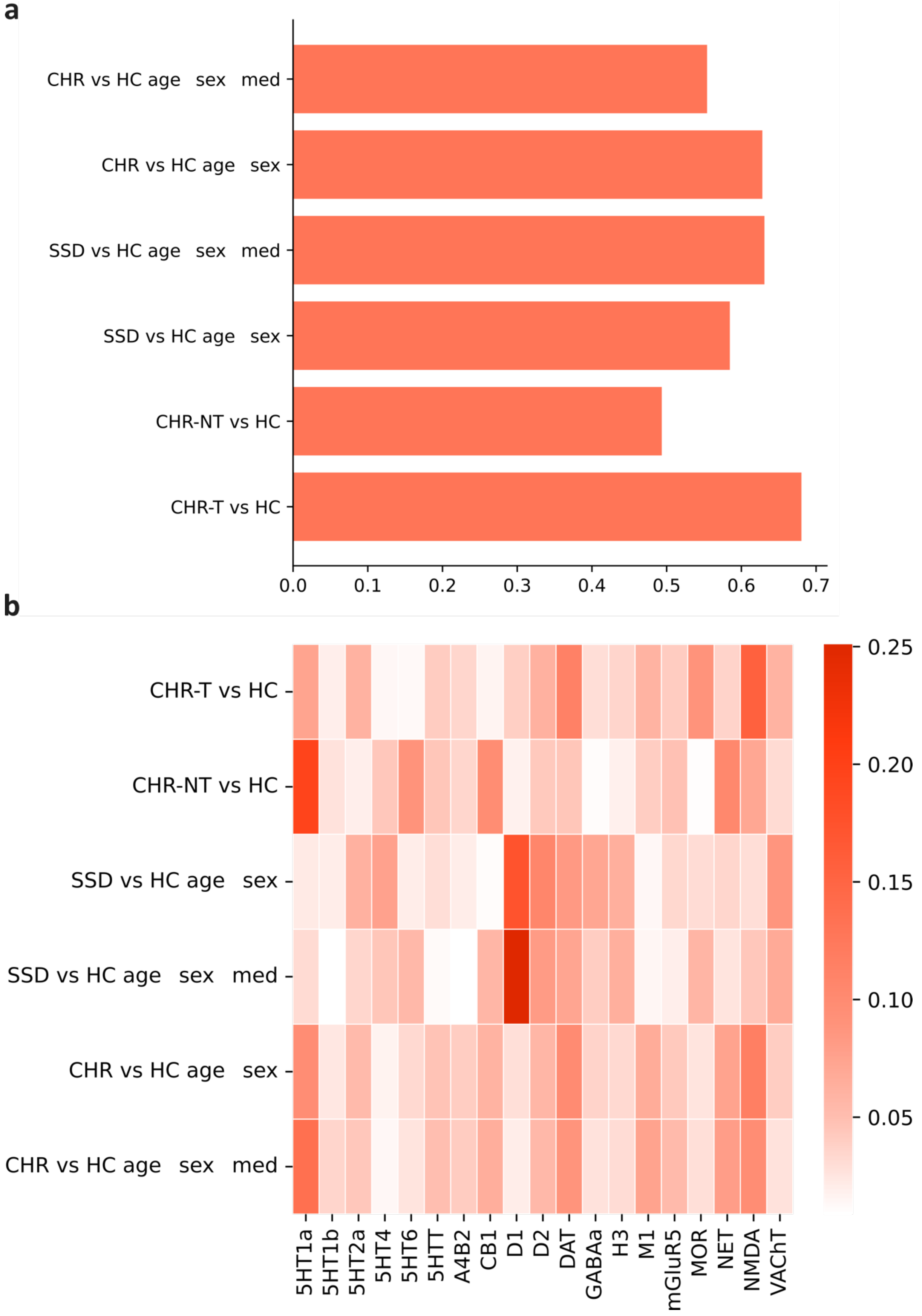

**Supplementary Fig. 5.** Dominance analysis of unique contribution of receptors to case-control cerebral blood flow (rCBF) phenotypes. **a**, Total dominance adjusted $R^2$. **b**, Individual total dominance, unique PET contribution to total adjusted $R^2$. 5HT1a/1b/2a/4/6, serotonin a/1b/2a/4/6 receptor. 5HTT, serotonin transporter. A4B2, alpha-beta-4-nicotinic receptor. CB1, Cannabinoid receptor 1. CHR, clinical high-risk for psychosis. D1/2, dopamine D1/2 receptor. DAT, dopamine transporter. GABAa, benzodiazepine binding site, γ-Aminobutyric acid A receptor. H3, histamine 3 receptor. M1, muscarinic receptor 1. mGluR5, Metabotropic glutamate receptor 5. MOR, µ-opioid receptor. NET, norepinephrine receptor. NMDA, *N*-methyl-D-aspartate receptor. SSD, schizophrenia spectrum disorder. VAChT, vesicular acetylcholine transporter.

**Dominance replication with DK atlas**

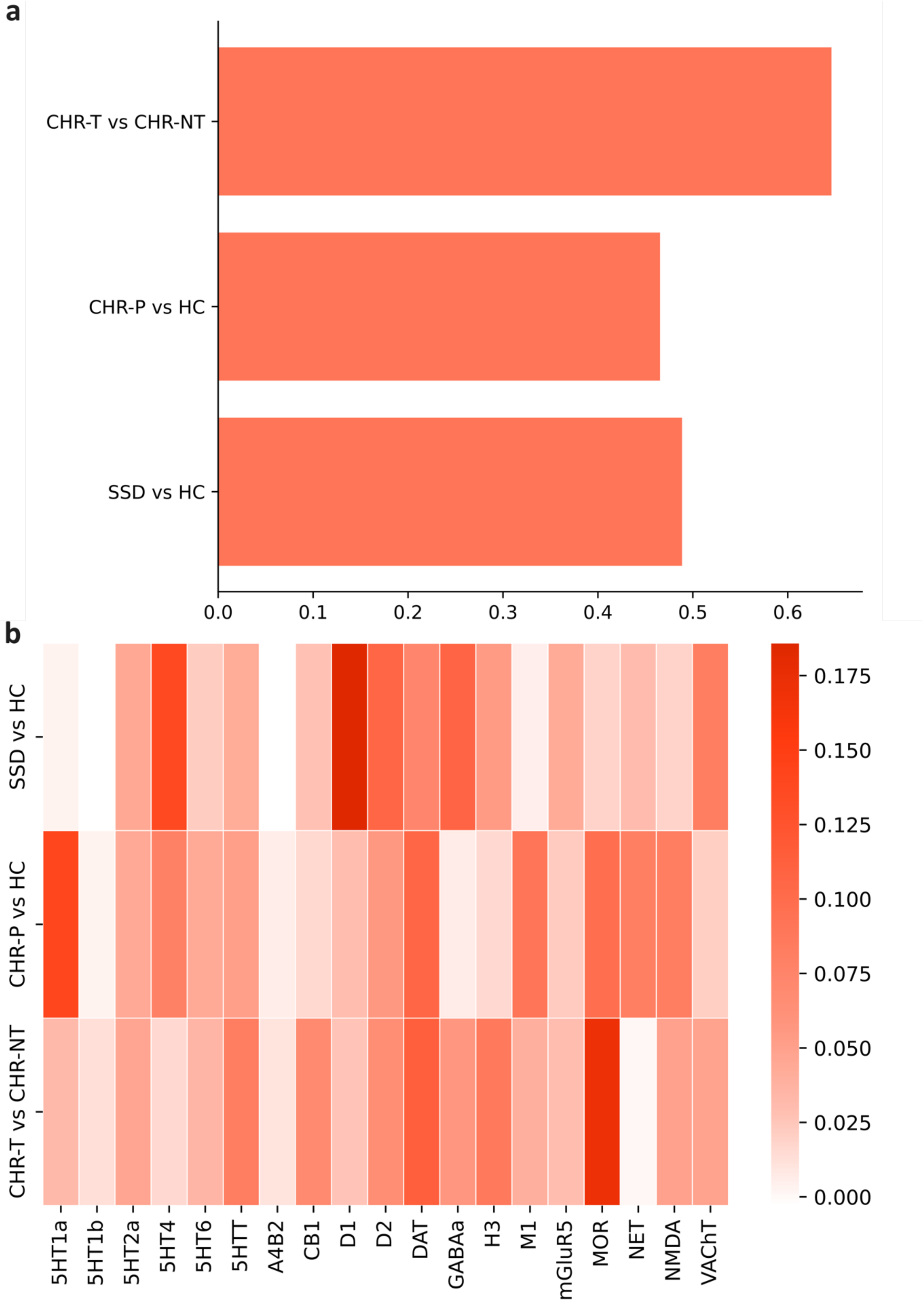

**Supplementary Fig. 6.** Replication of dominance analysis with DK-atlas, 82 ROIs. A) Total Dominance adjusted $R^2$. SSD vs HC adjusted $R^2$=.49, CHR-P vs HC adjusted $R^2$= .47, CHR-T adjusted $R^2$=.65, all models significant $p_{FDR}$<0.05. B) Individual predictor total dominance contribution. 5HT1a/1b/2a/4/6, serotonin a/1b/2a/4/6 receptor. 5HTT, serotonin transporter. A4B2, alpha-beta-4-nicotinic receptor. CB1, Cannabinoid receptor 1. D1/2, dopamine D1/2 receptor. DAT, dopamine transporter. GABAa, benzodiazepine binding site, γ-Aminobutyric acid A receptor. H3, histamine 3 receptor. M1, muscarinic receptor 1. mGluR5, Metabotropic glutamate receptor 5. MOR, μ-opioid receptor. NET, norepinephrine receptor. NMDA, *N*-methyl-D-aspartate receptor. VAChT, vesicular acetylcholine transporter.

**Dominance replication cortical only**

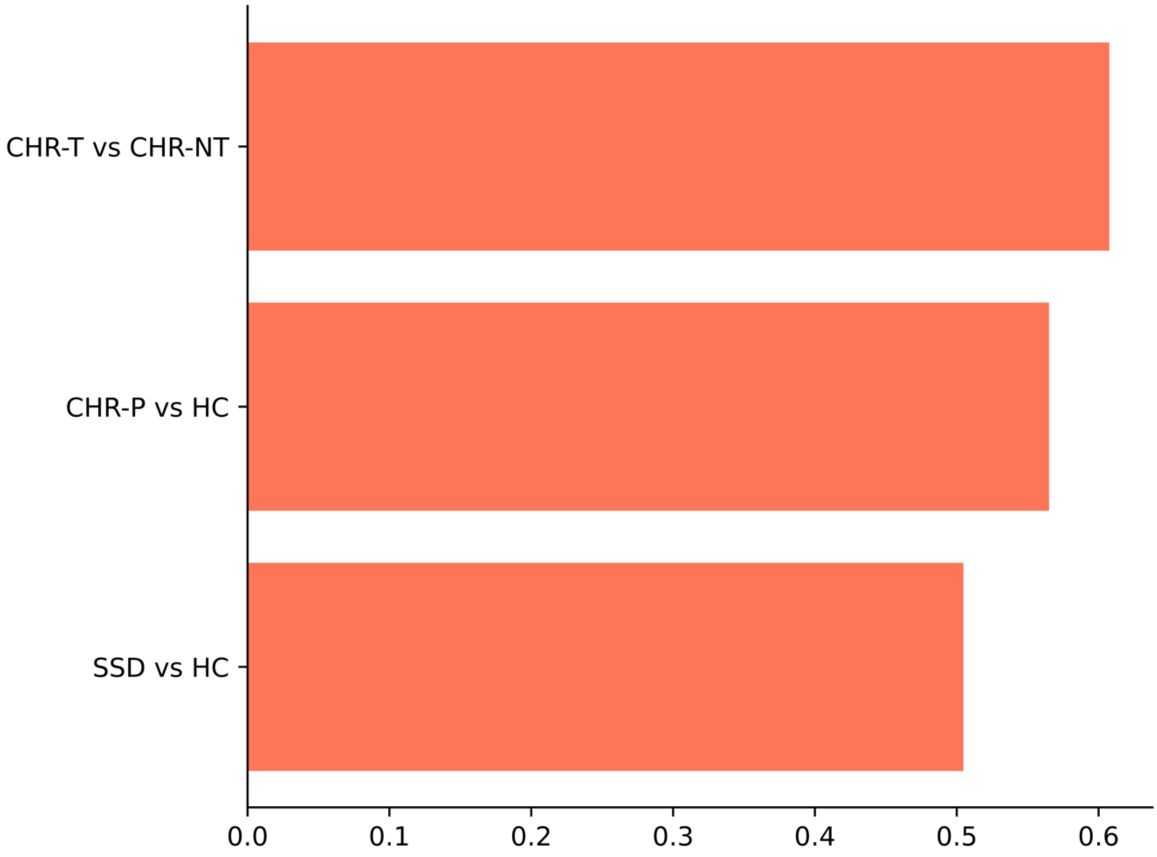

**Supplementary Fig. 7.** Replication of dominance analysis with Schaefer cortical atlas, 100 ROIs. No models were significant $p_{FDR}>0.05$.
